# Multivariate genome-wide association analysis of a cytokine network reveals variants with widespread immune, haematological and cardiometabolic pleiotropy

**DOI:** 10.1101/544445

**Authors:** Artika P. Nath, Scott C. Ritchie, Nastasiya F. Grinberg, Howard Ho-Fung Tang, Qin Qin Huang, Shu Mei Teo, Ari V. Ahola-Olli, Peter Würtz, Aki S. Havulinna, Kristiina Aalto, Niina Pitkänen, Terho Lehtimäki, Mika Kähönen, Leo-Pekka Lyytikäinen, Emma Raitoharju, Ilkka Seppälä, Antti-Pekka Sarin, Samuli Ripatti, Aarno Palotie, Markus Perola, Jorma S Viikari, Sirpa Jalkanen, Mikael Maksimow, Marko Salmi, Chris Wallace, Olli T. Raitakari, Veikko Salomaa, Gad Abraham, Johannes Kettunen, Michael Inouye

**Affiliations:** Cambridge Baker Systems Genomics Initiative, Baker Heart and Diabetes Institute, Melbourne, Victoria 3004, Australia; Cambridge Baker Systems Genomics Initiative, Department of Public Health and Primary Care, University of Cambridge, Cambridge CB1 8RN, United Kingdom; Department of Microbiology and Immunology, University of Melbourne, Parkville, Victoria 3010, Australia; Department of Medicine, University of Cambridge, Addenbrooke’s Hospital, Cambridge CB2 0SP, United Kingdom; Department of Clinical Pathology, University of Melbourne, Parkville, VIC 3010, Australia; Research Centre of Applied and Preventive Cardiovascular Medicine, University of Turku, Turku 20520, Finland; Satasairaala, Department of Internal Medicine, 28500 Pori, Finland; Institute for Molecular Medicine Finland, University of Helsinki, Helsinki 00014, Finland; Research Programs Unit, Diabetes and Obesity, University of Helsinki, Helsinki 00290, Finland; Nightingale Health Ltd, Helsinki 00300, Finland; National Institute of Health and Welfare, Helsinki 00271, Finland; Medicity Research Laboratory, Department of Medical Microbiology and Immunology, University of Turku, Turku 20520, Finland; Department of Clinical Chemistry, Fimlab Laboratories, and Finnish Cardiovascular Research Center - Tampere, Faculty of Medicine and Health Technology, Tampere University, Tampere 33520, Finland; Department of Clinical Physiology, Tampere University Hospital, and Finnish Cardiovascular Research Center - Tampere, Faculty of Medicine and Health Technology, Tampere University, Tampere 33521, Finland; Department of Public Health, University of Helsinki, Helsinki 00014, Finland; Broad Institute of MIT and Harvard, Cambridge, Massachusetts 02142, USA; Analytic and Translational Genetics Unit, Massachusetts General Hospital, Harvard Medical School, Boston, Massachusetts 02114, USA; Department of Psychiatry, Massachusetts General Hospital, Boston, Massachusetts 02114, USA; Department of Neurology, Massachusetts General Hospital, Boston, Massachusetts 02114, USA; Department of Medicine, University of Turku and Division of Medicine, Turku University Hospital, Turku 20520, Finland; Medicity Research Laboratory and Institute of Biomedicine, University of Turku, Turku 20520, Finland; MRC Biostatistics Unit, Institute of Public Health, Cambridge CB2 0SR, United Kingdom; The Department of Clinical Physiology and Nuclear Medicine, Turku University Hospital, Turku 20520, Finland; Computational Medicine, Centre for Life Course Health Research, University of Oulu, Oulu 90014, Finland; NMR Metabolomics Laboratory, School of Pharmacy, University of Eastern Finland, Kuopio 70211, Finland; Biocenter Oulu, University of Oulu 90014, Finland; The Alan Turing Institute, London, United Kingdom

## Abstract

Cytokines are essential regulatory components of the immune system and their aberrant levels have been linked to many disease states. Despite increasing evidence that cytokines operate in concert, many of the physiological interactions between cytokines, and the shared genetic architecture that underlie them, remain unknown. Here we aimed to identify and characterise genetic variants with pleiotropic effects on cytokines – to do this we performed a multivariate genome-wide association study on a correlation network of 11 circulating cytokines measured in 9,263 individuals. Meta-analysis identified a total of 8 loci significantly associated with the cytokine network, of which two (*PDGFRB* and *ABO*) had not been detected previously. Bayesian colocalisation analysis revealed shared causal variants between the eight cytokine loci and other traits; in particular, cytokine network variants at the *ABO, SERPINE2*, and *ZFPM2* loci showed pleiotropic effects on the production of immune-related proteins; on metabolic traits such as lipoprotein and lipid levels; on blood-cell related traits such as platelet count; and on disease traits such as coronary artery disease and type 2 diabetes.

## Introduction

Cytokines are signalling molecules secreted by cells that are central to multiple physiological functions, especially immune regulation (1). Broadly-speaking, cytokines include chemokines that drive movement of cells, and growth factors that drive cell growth and proliferation. Changes in circulating cytokine levels have been associated with infection (2), autoimmune diseases (3), malignancies (4), as well as atherosclerosis and cardiovascular disease (5,6). The expression of cytokines can be strongly regulated by genetic variation (7), and several studies have identified cis-acting genetic variants associated with circulating levels of certain cytokines and their receptors under various conditions (8–10). These initial studies laid the foundation for genetic investigation of circulating cytokine levels at a scale and breadth that may improve our understanding of individual differences in immune response, inflammation, infection and common disease susceptibility.

Despite cytokines operating in concert to facilitate immune regulation, genome-wide association studies (GWAS) have typically focused on individual cytokines (11–18). The most extensive cytokine GWAS to date separately analysed individual levels of 41 circulating cytokines in approximately 8,000 individuals, identifying 27 distinct loci each associated with at least one cytokine (19). Others have identified loci influencing cytokine production in response to pathogens (20,21). While these previous GWAS utilised a univariate framework, analysing each cytokine separately, studies of related traits indicate a multivariate framework can confer greater statistical power, for example by taking advantage of the tightly co-regulated nature of both pro and anti-inflammatory cytokines.

Several methods for multivariate GWAS of correlated phenotypes have been developed (22–27). Simulations have shown that multivariate analysis can result in increased power to detect genetic associations with small or pleiotropic effects across phenotypes (22,28–30). These have largely been conducted on metabolic traits where they have demonstrated a boost in statistical power. For example, multivariate analysis of four lipid traits led to a 21% increase in independent genome-wide significant variants compared to univariate analysis (23). Similar findings were shown for other metabolic traits (24,31). Moreover, complex genotype-phenotype dependencies have been revealed when jointly testing rare variants with lipoprotein traits (32). Notably, a multivariate GWAS of networks of highly correlated serum metabolites was able to detect nearly twice the number of loci compared to univariate testing, with downstream tissue-specific transcriptional analyses showing that the top candidate genes from multivariate analysis were upregulated in atherosclerotic plaques (31).

In this study, we focus on correlated immune traits by leveraging the correlation structure within a network of 11 cytokines to perform a multivariate genome-wide scan in 9,263 individuals from three population-based cohorts. We then investigate the colocalisation of cytokine-associated variants with those regulating gene expression in numerous tissues and cell types, circulating protein and metabolite levels, haematological traits, and disease states. Finally, we highlight and characterise variants as potential master regulator of the cytokine network, with pleiotropic effects on production of inflammatory proteins, immune cell function, lipoprotein and lipid levels, and cardiometabolic diseases.

## Methods

### Study populations

Approval for the study protocols for each cohort was obtained from their respective ethics committees, and all subjects enrolled in the study gave written informed consent.

The Cardiovascular Risk in Young Finns Study (YFS) is a longitudinal prospective cohort study commenced in 1980, with follow-up studies carried out every 3 years. The purpose of this study was to monitor the risk factors of cardiovascular disease in children and adolescents from different regions of Finland. In the baseline study, conducted in five Finnish metropolitan areas (Turku, Helsinki, Kuopio, Tampere and Oulu), a total of 3,596 children and adolescents were randomly selected from the national public register, the details of which were described in (33). A total of 2,204 participants responded to the 2007 follow-up study (YFS07), for which the age range was 30-45 years. Ethics were approved by the Joint Commission on Ethics of the Turku University and the Turku University Central Hospital.

The FINRISK cohorts were part of a cross-sectional population-based survey, which are carried out every five years since 1972 to evaluate the risk factors of chronic diseases in the Finnish population (34). Each survey has recruited a representative random sample of 6,000-8,800 individuals, within the age group of 25-74 years, chosen from the national population information system. This study utilised samples from the 1997 (FINRISK97) and 2002 (FINRISK02) collections, which recruited individuals from five or six (for FINRISK02) major regional and metropolitan areas of Finland: the provinces of North Karelia, Northern Savo, Northern Ostrobothnia, Kainuu, and Lapland; the Turku and Loimaa region of south-western Finland; and the Helsinki and Vantaa metropolitan area. In total, 8,444 (aged 24-74 years) and 8,798 (aged 51-74 years) individuals participated in the FINRISK97 and FINRISK02 studies, respectively. Importantly, each FINRISK survey is an independent cohort, each comprising a different set of participants. Ethics were approved by the coordinating ethical committee of the Helsinki and Uusimaa hospital district, Finland.

### Blood sample collection

Blood samples and detailed information on various physical and clinical variables for the YFS and FINRISK cohorts were collected using similar protocols as described previously (33,34). Venous blood was collected following an overnight fast for the YFS cohorts, while non-fasting blood was collected for FINRISK. Samples were centrifuged, and the resulting plasma and serum samples were aliquoted into separate tubes and stored at −70°C for later analyses.

### Genotype processing and quality control

Genotyping in YFS and FINRISK cohorts was performed on whole blood genomic DNA. For YFS07 (N=2,442), a custom 670K Illumina BeadChip array was used for genotyping. For FINRISK97 (N=5,798), the Human670-QuadCustom Illumina BeadChip platform was used for genotyping. For FINRISK02 (N=5,988), the Human670-QuadCustom Illumina BeadChip (N=2,447) and the Illumina Human CoreExome BeadChip (N=3,541) was used for genotyping. The Illuminus clustering algorithm was used for genotype calling (35) and quality control (QC) was performed using the Sanger genotyping QC pipeline. This included removing SNPs and samples with > 5% genotype missingness followed by removal of samples with gender discrepancies. Genotypes were then imputed with IMPUTE2 (36) using the 1000 Genomes Phase 1 version 3 as the reference panel followed by removal of SNPs with call rate < 95%, imputation “info” score < 0.4, minor allele frequency < 1%, and Hardy-Weinberg equilibrium *P*-value < 5 × 10^−6^. Instances where data was generated using different genotyping platforms, overlapping SNPs were merged using PLINK version 1.90 software (https://www.cog-genomics.org/plink2) (37). A total of 6,664,959, 7,370,592 and 6,639,681 genotyped and imputed SNPs passed quality control in YFS, FINRISK97 and FINRISK02, respectively. Cryptic relatedness was assessed using identity by descent (IBD) estimates and in cases where the pi-hat relatedness was greater than 0.1, one of the two individuals was randomly removed (N=44 for YFS, N=291 for FINRISK97, and N=39 for FINRISK02). Genetic PCs were obtained through principal component analysis (PCA) using FlashPCA (38) on ~60,000 LD pruned SNPs.

### Measurement of cytokines

Concentrations of cytokines, chemokines, and growth factors (hereafter referred to as cytokines) were measured in serum (YFS07), EDTA plasma (FINRISK97), and heparin plasma (FINRISK02) using multiplex fluorescent bead-based immunoassays (Bio-Rad). A total of 48 cytokines were measured in YFS07 (N=2,200) and FINRSK02 (N=2,775) using two complementary array systems: the Bio-Plex ProTM Human Cytokine 27-plex assay and Bio-Plex ProTM Human Cytokine 21-plex assay. For FINRISK97, 19 cytokines were assayed on the Human Cytokine 21-plex assay system. All assays were performed in accordance with the manufacturer’s instructions, except that the amount of beads, detection antibodies, and streptavidin-phycoerythrin conjugate were used at half their recommended concentration. Fluorescence intensity values determined using the Bio-Rad’s Bio-Plex 200 array reader were converted to concentrations from the standard curve generated by the Bio-Plex^™^ Manager 6.0 software. For each cytokine, a standard curve was derived by fitting a five-parameter logistic regression model to the curve obtained from standards provided by the manufacturer. Cytokines with concentrations at the lower and upper asymptotes of the sigmoidal standard curve were set to the concentration corresponding to the fluorescent intensity 2% above or below the respective asymptotes.

### Cytokine data filtering, normalisation and clustering

The analysis was limited to 18 cytokines (**Table S1**) assayed in all three cohorts. Although Interleukin 1 receptor, type I (IL-1Ra) was assayed in all three cohorts, it was excluded from the analyses due to its inconsistent Pearson correlation pattern with other 18 cytokines across the three datasets.

Before normalisation, cytokine data was subset to individuals with matched genotype data in YFS07 (N=2,018), FINRISK97 (N=5,728), and FINRISK02 (N=2,775). We excluded individuals in YFS07 reporting febrile infection in the two weeks prior to blood sampling (N=92). To identify extreme outlier samples, PCA was performed on the log2 transformed cytokine values using the missMDA R package (39). This method first imputed the missing cytokine values using a regularised iterative PCA algorithm implemented in the imputePCA function, before performing PCA. Three and two outlier samples were removed from FINRISK97 and FINRISK02 respectively. Based on IBD analysis described above, 44 (YFS07), 291 (FINRISK97), and 39 (FINRISK02) individuals were also removed. After filtering, a total of 1,843, 5,434 and 1,986 individuals passed quality control in YFS07, FINRISK97 and FINRISK02, respectively, and these were used for downstream analysis.

Since all 18 cytokines displayed non-Gaussian distributions, we performed normalisation of cytokine levels. For YFS07, the lower limit of detection (LOD) was available for each cytokine. Reported values that were below the LOD were indistinguishable from background noise signals or instrument error (40), and were excluded and treated as missing. For FINRISK97 and FINRISK02, the detection limits were not available; however, it was observed that these two datasets exhibited a bimodal distribution, with the leftmost peak below the expected LOD when compared to the YFS dataset. Individuals in the leftmost peak were therefore set to missing. The log2-transformed cytokine values were then normalised to follow standard Gaussian distributions (with mean of 0 and sd of 1) using rank-based inverse normal transformation (rntransform) as implemented in the GenABEL R package (41). For each study group, residuals for all cytokines were calculated by regressing the normalised cytokine values on age, sex, BMI, lipid and blood pressure medication, pregnancy status (FINRISK97), and the first 10 genetic PCs using a multiple linear regression model.

Detection of groups of correlated cytokines was done in FINRISK97, the cohort with the largest sample size. Pairwise Pearson correlation was performed amongst residuals of 18 cytokines. These cytokines were then subjected to hierarchical clustering, with one minus the absolute correlation coefficient used as the dissimilarity metric. We then defined a cytokine network - a group of 11 cytokines that were moderate-to highly-correlated (r > 0.57) - for subsequent use in the multivariate analysis.

### Statistical Analysis

Univariate association analysis was carried out with linear regression in PLINK (37), where the residuals of each cytokine were regressed on each SNP genotypes. Summary statistics at each marker across three datasets were then combined in a meta-analysis using the METAL software program (42), which implemented a weighted Z-score method.

Multivariate testing (MV) was performed under the canonical correlation framework implemented in PLINK (MV-PLINK) (22), which extracted the linear combination of traits most highly-correlated with genotypes at a particular SNP. The test is based on Wilks’ Lambda (λ = 1−ρ^2^), where ρ is the canonical correlation coefficient between the SNP and the cytokine network. Corresponding *P*-values were computed by transforming Wilks’ Lambda to a statistic that approximates an *F* distribution and the loadings for each cytokine represented their individual contributions toward the multivariate association result (22). Since the multivariate beta-coefficients and standard errors were not calculated by MV-PLINK, the cohort-level multivariate *P*-values were combined in a meta-analysis using the weighted Z-score method (43,44) implemented in the metap R package. Briefly, the *P*-values for each dataset were transformed into Z-scores, weighted by their respective sample sizes and the sum of these weighted Z-scores were then divided by the square root of the sum of squares of the sample size for each study. The combined weighted Z-score obtained was back-transformed into a one-tailed *P-*value.

To assess the inflation of the test statistics as a result of population structure, quantile-quantile (Q-Q) plots of observed *vs.* expected-log_10_ *P*-values were generated from the multivariate analysis of the three datasets, both individually and meta-analysed. Corresponding genomic inflation factor (λ) was calculated by taking the ratio of the median observed distribution of *P*-values to the expected median.

To investigate the existence of additional independent signals within the significant multivariate loci, a conditional stepwise multivariate meta-analysis was performed within each locus. For each study cohort, the lead SNP at each locus (*P*-value < 5 × 10^−8^) together with other covariates were fitted in a linear regression model for each cytokine in the network. The resulting residuals were provided as an input for the multivariate test of the locus being assessed. The cohort-level conditional *P*-values were then combined in a meta-analysis. The stepwise conditional analysis was repeated in the univariate model with the lead multivariate SNPs until no additional significant signal was identified.

### Colocalisation analysis

Bayesian colocalisation tests between cytokine network-associated signals and the following trait- and disease-associated signals were performed using the COLOC R package (45). For whole blood *cis*-eQTLs, we downloaded publicly-available summary data from the eQTLGen Consortium portal (http://www.eqtlgen.org/). The eQTLGen Consortium analysis is the largest meta-analysis of blood eQTLs to date and comprises of 31,684 blood and PBMC samples from a total of 37 datasets (46). For immune cell *cis*-eQTLs, we either generated *cis*-eQTL summary data in resting B-cells (47), resting monocytes (48), and stimulated monocytes with interferon-γ or lipopolysaccharide (48), or obtained publicly-available *cis*-eQTL summary data generated by the BLUEPRINT consortium in neutrophils and CD4^+^ T-cells (49). For *cis*-eQTL mapping in B-cells and monocytes (resting and stimulated), information on accessing the raw gene expression and genotype data, data pre-processing, and *cis*-eQTL analysis has been described in a previous study (50). The BLUEPRINT immune cell summary statistics was downloaded from: ftp://ftp.ebi.ac.uk/pub/databases/blueprint/blueprint_Epivar/. For protein QTLs, we used publicly-available SomaLogic plasma protein GWAS summary statistics from the INTERVAL study (17). For disease or complex trait associations, we compiled summary statistics of 185 diseases and quantitative traits from GWAS studies conducted in European ancestry individuals, which were accessed from the UK biobank (**Table S10**), or downloaded from either ImmunoBase (https://www.immunobase.org/), the NHGRI-EBI GWAS Catalog (https://www.ebi.ac.uk/gwas/), or LD Hub (http://ldsc.broadinstitute.org/). Here, we only considered immune-related and cardiometabolic diseases. For each cytokine network locus, we only tested traits or diseases with the minimum association *P*-value < 1 × 10^−6^ at this locus. COLOC requires either beta-coefficients and its variance, or *P*-values, for each SNP, in addition to MAF and sample size. Since PLINK multivariate did not produce beta values and standard errors, we instead used meta-analysed *P-*values for the multivariate cytokine GWAS summary data. For each association pair assessed for colocalisation, SNPs within 200kb of the lead multivariate cytokine GWAS SNP were considered. COLOC (coloc.abf) was run with default parameters and priors. COLOC computed posterior probabilities for the following five hypotheses: PP0, no association with trait 1 (cytokine GWAS signal) or trait 2 (e.g. eQTL signal); PP1, association with trait 1 only (i.e. no association with trait 2); PP2, association with trait 2 only (i.e. no association with trait 1); PP3, association with trait 1 and trait 2 by two independent signals; PP4, association with trait 1 and trait 2 by shared variants. In practice, evidence of colocalisation were defined by PP3 + PP4 ≥ 0.99 and PP4/PP3 ≥ 5, a cut off previously suggested (50).

## Results

### Summary of cohorts and data

Our final dataset comprised a total of 9,267 individuals enrolled in three population-based studies, YFS07 (N=1,843), FINRISK97 (N=5,438), and FINRISK02 (N=1,986), all of whom had genome-wide genotype data and quantitative measurements of 18 cytokines (**Table S1**). Characteristics of the study cohorts are summarised in **Table 1**. Genotypes for the three datasets were imputed with IMPUTE2 (36) using the 1000 Genomes Phase 1 version 3 of the reference panel. After quality control, a total of 6,022,229 imputed and genotyped SNPs were available across all cohorts. Cytokine levels were measured in serum and plasma using Bio-Plex ProTM Human Cytokine 27-plex and 21-plex assays, then subsequently normalised and adjusted for covariates including age, sex, BMI, pregnancy status, blood pressure lowering medication, lipid lowering medication, and population structure (**Methods**). An overview of the study is shown in **Figure 1**.

**Table 1:**
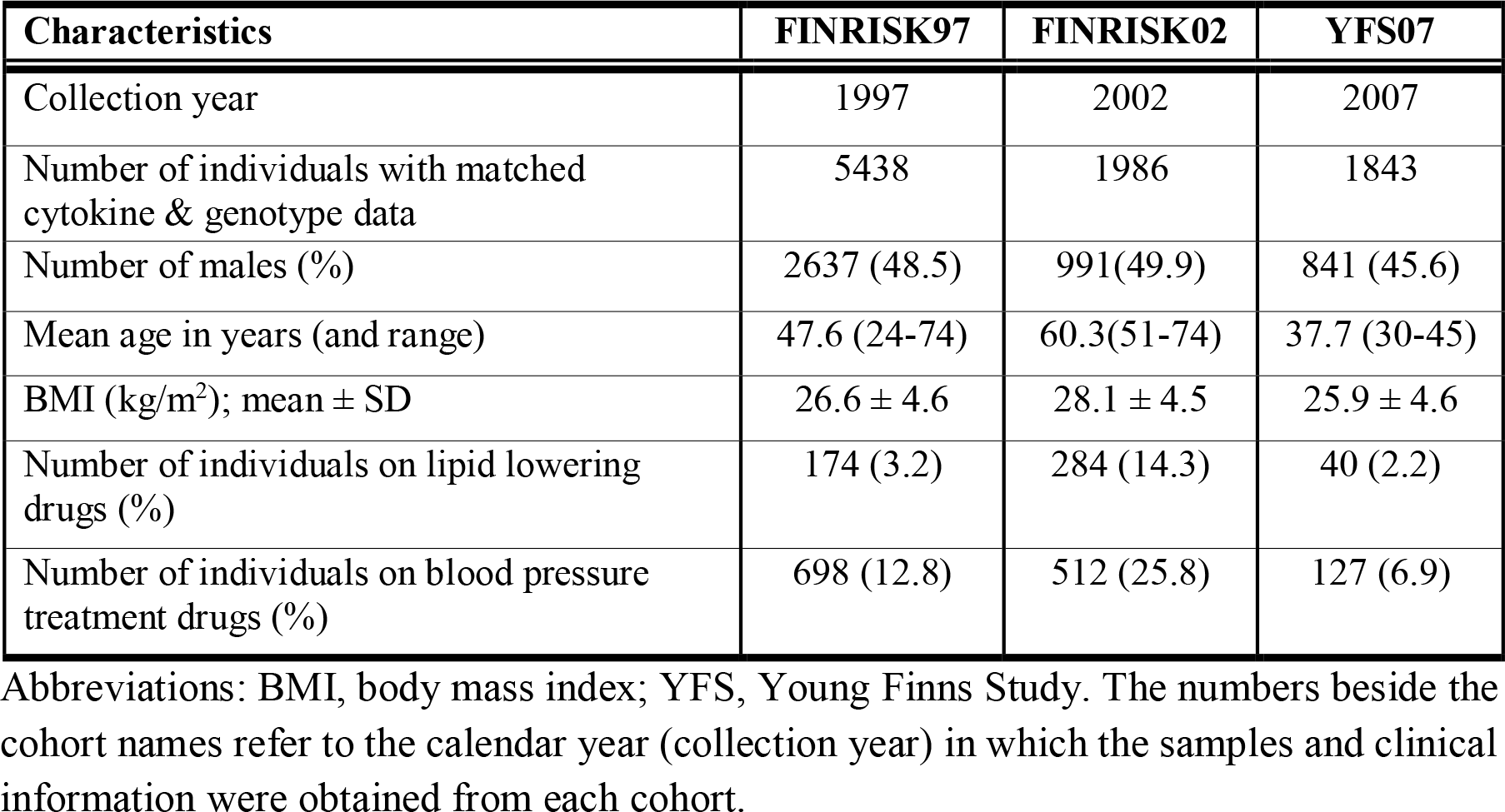
Summary of descriptive characteristics of the three study cohorts.

**Figure 1:**
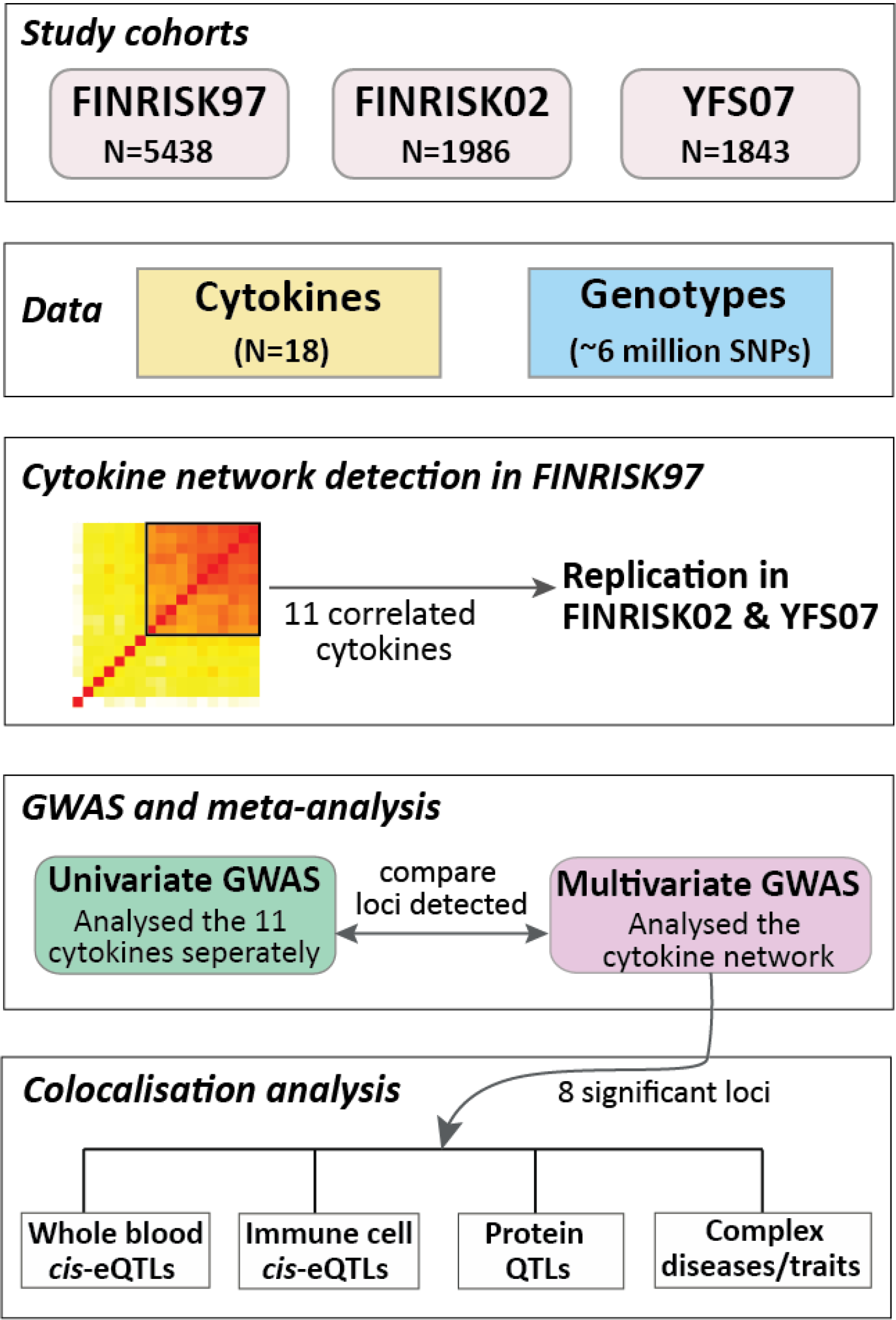
Overview of the study populations, design, and the analyses conducted.

### A correlation network of circulating cytokines

To characterise the correlation structure of circulating cytokines, we utilised the largest dataset available (FINRISK97) and the set of 18 cytokines overlapping all three cohorts. IL-18 was very weakly correlated with other cytokines (**Figure 2A**), while TRAIL, SCF, HGF, MCP-1, EOTAXIN and MIP-1b showed moderate correlation with the others. A distinct set of 11 cytokines showed high correlation amongst themselves (median r=0.75). In the smaller cohorts (YFS07 and FINRISK02), the cytokine correlation structure was similar but weaker (**Figure S1**), with the set of 11 cytokines also showing relatively high correlation (YFS07 median r=0.42; FINRISK02 median r=0.46). We utilised this set of 11 cytokines (denoted below as the cytokine network) for multivariate association analysis.

**Figure 2:**
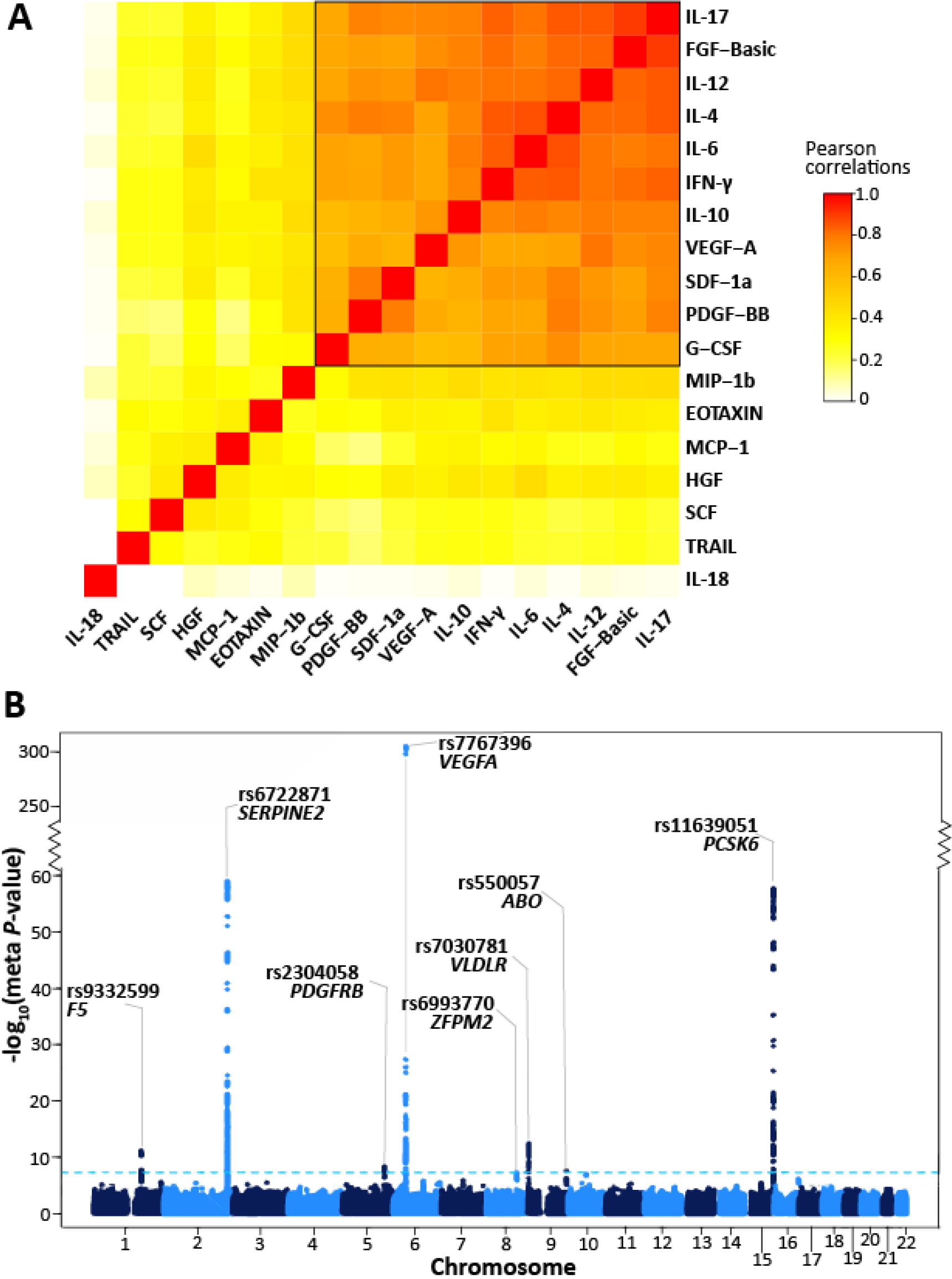
Multivariate GWA analysis of a network of 11 correlated cytokines in three Finnish cohorts. **(A)** Correlation heatmap of the 18 cytokines in the FINRISK97 cohort. Each cell presents the pair-wise Pearson’s correlation coefficient between the normalised cytokine residuals. The cytokines are ordered by hierarchical clustering, using 1 minus the absolute value of the correlations as the distance matrix. The colour scale denotes the strength of the correlations, where red is a high positive correlation. The group of 11 tightly correlated cytokines (black box) was used for multivariate analysis. (**B)** Manhattan plot for meta-analysis results from the multivariate GWAS of the cytokine network. The statistical strength of association (−log_10_ meta-*P*-value; *y*-axis) is plotted against all the SNPs ordered by chromosomal position (*x*-axis). The sky-blue horizontal dashed line represents the genome-wide (meta-*P*-value < 5 × 10^−8^) significance threshold. The lead SNP (lowest meta-*P*-value) at each locus and the nearby genes are shown.

The cytokine network included both anti-inflammatory (IL-10, IL-4, IL-6) and pro-inflammatory (IL-12, IFN-γ, IL-17) cytokines as well as growth factors (FGF-basic, PDGF-BB, VEGF-A, G-CSF) and a chemokine (SDF-1a) involved in promoting leukocyte extravasation and wound healing (51–53). These cytokines were all positively correlated, which is likely indicative of counter-regulatory (negative-feedback) mechanisms amongst pro-inflammatory and anti-inflammatory pathways, such as that of IFN-γ and IL-10 (54).

### Multivariate genome-wide association analysis for cytokine loci

We performed a multivariate GWAS on the cytokine network in each cohort separately, then cohort-level results were combined using meta-analysis (**Methods**). Since one hypothesis test (corresponding to the cytokine network) was performed for each SNP, a genome-wide significance threshold of *P* < 5 × 10^−8^ was used. Minimal inflation was observed for the cohort-level and meta-analysis test statistics with lambda (λ) inflation ranging between 1.00-1.02 (**Figure S2A - D**).

We identified 8 loci reaching genome-wide significance for the cytokine network (**Figure 2B; Table 2**). The strongest association was rs7767396 (meta-*P-*value = 6.93 × 10^−306^), a SNP located 172kb downstream of vascular endothelial growth factor A (*VEGFA*) (**Figure S3A**). The *VEGFA* locus was previously identified in GWAS for individual cytokine levels including VEGF-A, IL-7, IL-10, IL-12, and IL-13 (14,19). Consistent with these earlier results, we found that VEGF-A, IL-10, and IL-12 were the top three cytokines based on their trait loadings (relative contribution of each cytokine to the multivariate association result) in each cohort and also significantly associated with this locus in the univariate scans (**Figure S4A**). Multivariate analysis also confirmed four other previously known associations (14,16,19), including loci harbouring *SERPINE2* (rs6722871; meta-*P-*value = 1.19 ×10^−59^), *ZFPM2* (rs6993770; meta-*P-* value = 4.73 × 10^−8^), *VLDLR* (rs7030781; meta-*P-*value = 3.78 × 10^−13^), and *PCSK6* (rs11639051; meta-*P-*value = 1.93 × 10^−58^) (**Figure 2B; Table 2; Figure S3B - E**). The cytokine with the highest loading at each of these loci was consistent with those previously identified in univariate analysis (**Figure S4B - E**).

**Table 2:** Meta-analysed results of multivariate GWAS of cytokine network.

| Locus | Locus Region | Top SNP | Average MAF | Top Multivariate Meta-*P*-value | Univariate Meta-*P*-value (Top Cytokine) | Detection |
| --- | --- | --- | --- | --- | --- | --- |
| *F5* | 1q24.2 | rs9332599 | 0.294 | $7.17 \times 10^{-12}$ | $9.21 \times 10^{-3}$ (SDF1a) | Multivariate |
| *SERPINE2* | 2q36.1 | rs6722871 | 0.311 | $1.19 \times 10^{-59}$ | $3.55 \times 10^{-18}$ (PDGF-BB) | Both |
| *PDGFRB* | 5q32 | rs2304058 | 0.379 | $4.06 \times 10^{-9}$ | $1.52 \times 10^{-5}$ (IL4) | Multivariate |
| *VEGFA* | 6p21.1 | rs7767396 | 0.471 | $6.93 \times 10^{-306}$ | $3.10 \times 10^{-201}$ (VEGF-A) | Both |
| *ZFPM2* | 8q23.1 | rs6993770 | 0.221 | $4.73 \times 10^{-8}$ | $1.01 \times 10^{-7}$ (IL12p70) | Multivariate |
| *ABO* | 9q34.2 | rs550057 | 0.306 | $2.75 \times 10^{-8}$ | $4.9 \times 10^{-3}$ (IL4) | Multivariate |
| *VLDLR* | 9p24.2 | rs7030781 | 0.413 | $3.78 \times 10^{-13}$ | $6.78 \times 10^{-14}$ (VEGF-A) | Both |
| *PCSK6* | 15q26.3 | rs11639051 | 0.255 | $1.93 \times 10^{-58}$ | $1.19 \times 10^{-26}$ (PDGF-BB) | Both |
| *JMJD1C* | 10q21.3 | rs9787438 | 0.374 | *$1.30 \times 10^{-7}$ | *$8.96 \times 10^{-12}$ (VEGFA) | Univariate |

The table shows the meta-analysis *P*-values for the top SNP (lowest *P*-value) at each locus associated with the cytokine network in the multivariate analysis at genome-wide significance threshold ( $5 \times 10^{-8}$ ). The corresponding lowest meta-*P*-value for the same top SNP in the univariate analysis with any single cytokine present in the cytokine network, given in brackets beside the meta-*P*-value, was also reported. \*Instance where the top SNP at a locus crossed only the univariate significance threshold ( $P < 4.55 \times 10^{-9}$ ), then the corresponding meta-*P*-value for that SNP in the multivariate was also given. The univariate significance threshold was calculated from a Bonferroni correction for 11 cytokines tested ( $5 \times 10^{-8}/11$ ).

The multivariate GWAS also detected novel cytokine associations not identified in any previous univariate tests of these cytokines. These were three loci with genic lead SNPs in the candidate genes *F5, PDGFRB*, and *ABO*. The lead variant at the *F5* locus (rs9332599; meta-*P-*value = 7.17 ×10^−12^) is located in intron 12 of *F5* (**Figure S3F**). At the platelet-derived growth factor receptor-beta (*PDGFRB*) locus, the lead variant rs2304058 (meta-*P-*value = 4.06 × 10^−9^) is within intron 10 of *PDGFRB* (**Figure S3G**). At the *ABO* locus, the lead variant rs550057 (meta-*P-*value = 2.75 × 10^−8^) is within the first intron of *ABO* (**Figure S3H**); furthermore, rs550057 is located ~1.6 kb upstream of the erythroid cell specific enhancer, which contains a GATA-1 transcription factor binding site and has been shown to enhance the transcription of the *ABO* gene (55).

To investigate the presence of multiple independently associated variants at each of the eight loci, we performed stepwise conditional multivariate meta-analysis. Three loci (*SERPINE2*, *VEGFA*, and *PCSK6*) exhibited evidence of multiple independent signals (**Table S2**). In addition to the lead variants (rs6722871, rs7767396, rs11639051) at each of these three loci, we identified additional association signals (rs55864163; *SERPINE2*, meta-*P_cond._* = 9.03 × 10^−29^; rs112215592, *SERPINE2*, meta-*P_cond_* = 2.10 × 10^−12^; rs4714729; *VEGFA*, meta-*P_cond_* = 7.49 × 10^−10^; rs6598475, *PCSK6*, meta-*P_cond_* = 2.63 × 10^−17^), which were independently associated with the cytokine network. We also performed conditional univariate analysis that adjusted for the lead multivariate SNPs, which were either the same lead univariate SNPs or in high LD (r^2^ = 0.99). This univariate analysis also uncovered the same secondary signal at the *VEGFA* locus in association with VEGFA cytokine levels (rs4714729; meta-*P_cond_* = 8.8 × 10^−13^) (**Table S2**).

### Colocalisation of cytokine variants with cis-eQTLs in whole blood

To characterise the regulatory effects of the multivariate cytokine-associated loci, we queried the largest publicly-available set of results for whole blood *cis*-eQTLs from a meta-analysis of 31,684 individuals, which was obtained from the eQTLGen Consortium database (46). We found SNPs, lead or LD-proxy (r^2^>0.5), at seven of the eight cytokine loci (*ABO*, *F5*, *PCSK6*, *PDGFRB*, *SERPINE2*, *VEGFA*, *VLDLR*) with *cis*-regulatory effects (*P*-value < 1 × 10^−6^) on gene expression (a total of 17 unique genes) in blood (**Table S3)**. Using Bayesian colocalisation analysis, we further demonstrated that associations at three of these loci colocalised with *cis*-eQTLs for *ABO*, *PCSK6*, and *SERPINE2* expression (**Figure 3A - C; Table S4**).

**Figure 3:**
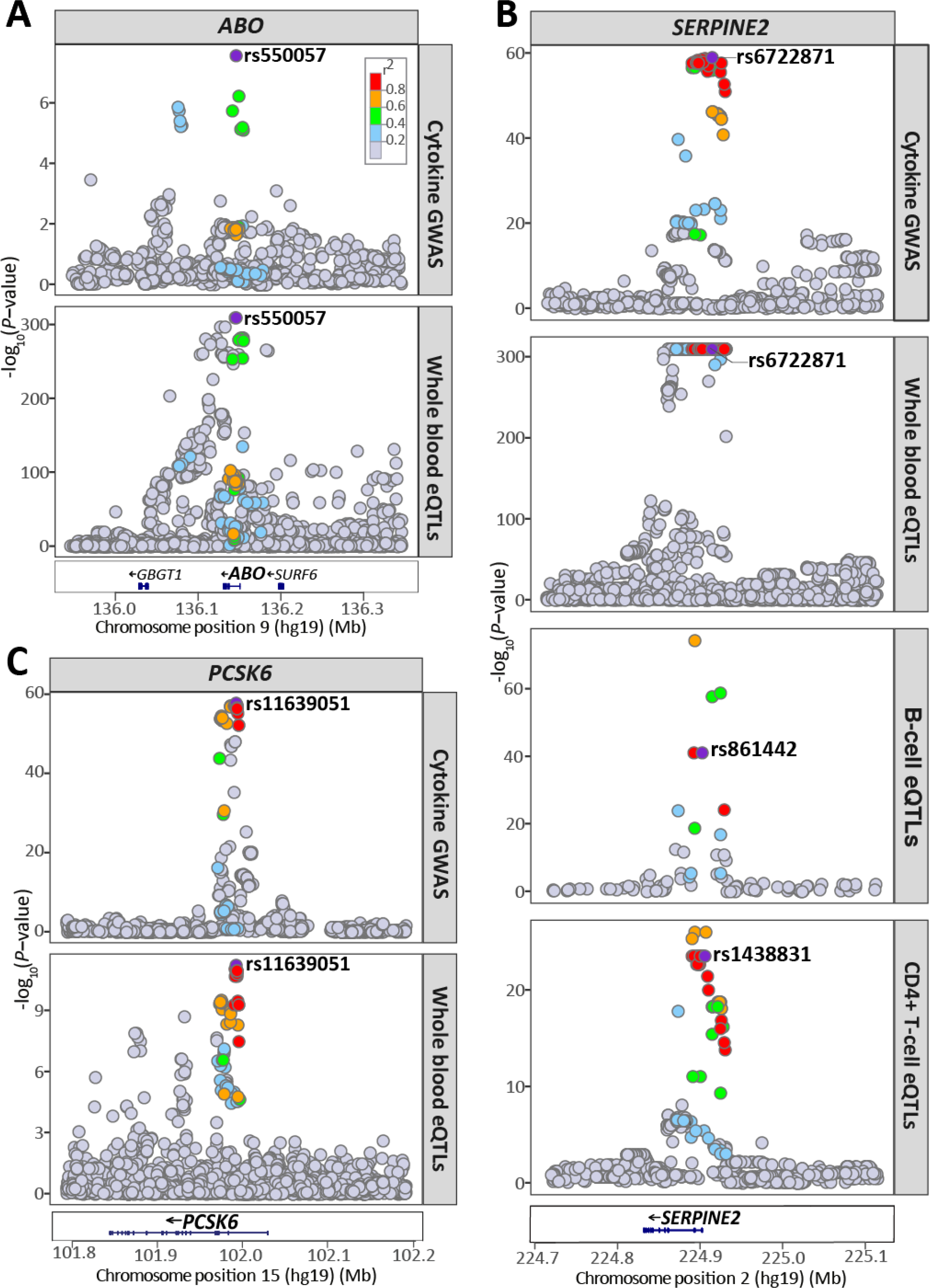
Regional plots for the cytokine network association, and whole blood and immune cell *cis*-eQTL association signals at the *ABO*, *PCSK6* and *SERPINE2* locus. **(A)** The cytokine network GWAS signal (top) colocalises with the whole blood *cis*-eQTLs signal for *ABO* (bottom) at the *ABO* locus on chromsome 9; **(B)** colocalises with whole blood *cis*-eQTLs for *PCSK6* expression (bottom) at the *PCSK6* locus on chromosome 15; **(C)** colocalises with the *cis*-eQTL signals for *SERPINE2* expression in whole blood (middle), B-cells (middle), and CD4^+^ T-cells (bottom) at the *SERPINE2* locus on chromosome 2. For each plot, the circles represent the −log_10_ association *P*-values (*y*-axis) of SNPs plotted against their chromosomal position (*x*-axis). The eQTL association plots show the lead cytokine network GWAS SNP tested in the colocalisation analysis. The lead cytokine network GWAS SNP rs6722871 was not present in the B-cell and CD4^+^ T cell eQTL dataset, instead, the next top GWAS SNP present in each of the eQTL dataset (rs861442, B-cell; rs1438831, CD4^+^ T-cell) is shown. For all regional plots, pairwise LD (r^2^) in the region is coloured with respect to the lead cytokine network GWAS SNP. LD was calculated from the 1000 Genomes European population.

### Colocalisation of cytokine variants with immune cell-specific cis-eQTLs

Next, we investigated the cell type- or context-dependent regulatory effects of genetic variants associated with the cytokine network by interrogating previously published *cis*-eQTLs specific to resting B-cells (47), resting monocytes (48), stimulated monocytes with interferon-γ or lipopolysaccharide (48), resting neutrophils (56), naive CD4^+^ T-cells (49,56) and CD8^+^ T-cells (49), all isolated from healthy donors of European ancestry (**Table S5**). Three out of the eight cytokine network loci harboured *cis*-eQTLs (*P*-value < 1 × 10^−6^) in at least one immune cell type, in either stimulated or non-stimulated state (**Table S6**). For example, SNPs at the *SERPINE2* locus were reported to have *cis*-eQTL effects across multiple immune cell types, including B-cells, CD4^+^ and CD8^+^ T-cells (**Table S6**).

Further, colocalisation analysis showed that the cytokine network variants at *SERPINE2* had strong evidence of sharing a causal variant with *SERPINE2 cis*-eQTLs in CD4^+^ T-cells and B-cells, similar to the colocalisation we observe in whole blood (**Figure 3B**; **Table S7**).

### Colocalisation of cytokine variants with plasma protein QTLs

To investigate protein-level effects of cytokine network variants, we utilised plasma protein QTLs (pQTLs) from the INTERVAL study (17). Colocalisation analysis, considering only pQTLs with association *P*-value < 1 × 10^−6^, showed all the eight cytokine network loci had strong evidence of shared causal variants with plasma levels of a total of 146 proteins (out of the 215 tested) (**Table S8**). Of these, the *ABO* and *ZFPM* cytokine network loci strongly colocalised with pQTL signals for 55 (out of 81) and 87 (out of 98) proteins, respectively (**Table 3; Table S8**). Of these, 14 and 75 proteins shared the same causal lead pQTLs with the lead cytokine network variants at the *ABO* (rs550057) and *ZFPM2* (rs6993770) loci, respectively, suggesting these variants have widespread effects.

**Table 3:** Colocalisation of cytokine network-associated variants at the *ABO* and *ZFPM2* loci with those of plasma protein levels, quantitative traits, and disease risk. Evidence: evidence of colocalisation; Strong: PP3+PP4 > 0.99 and PP4/PP3 > 5; Suggestive: PP3 + PP4 > 0.75 and PP4/PP3 > 3; None: association signal for the trait at the locus, but no evidence of colocalisation.

| ABO locus (Chromosome 9) |  |  |  |
| --- | --- | --- | --- |
| Traits/<br>Diseases | Group/ Functions | Evidence | Names |
| Diseases | Cardiometabolic diseases | Strong | Pulmonary embolism, ischemic stroke, coronary artery disease, type 2 disease, |
|  |  | None | Deep vein thrombosis |
| Blood cell traits | Blood cell counts | Strong | White blood cell, granulocytes, basophils + eosinophils, basophils + neutrophils, eosinophils + neutrophils, eosinophils, neutrophils, haematocrit (%), haemoglobin, myeloid, red blood cells, platelet distribution width |
|  |  | Suggestive | Basophils, reticulocytes |
|  |  | None | Monocyte, platelet, plateletcrit (%), red cell distribution width |
| Metabolites | IDL particle constituents | Strong | Total cholesterol (IDL-C), free cholesterol (IDL-FC), total lipids (IDL-L), total particle concentration (IDL-P), phospholipids (IDL-PL), triglycerides (IDL-TG) |
|  | LDL subclass particle constituents | Strong | For large particles: total cholesterol (L-LDL-C), cholesterol esters (L-LDL-CE), free cholesterol (L-LDL-FC), total lipids (L-LDL-L), total particle concentration (L-LDL-P), phospholipids (L-LDL-PL),<br>For medium particles: total cholesterol (M-LDL-C), cholesterol esters (M-LDL-CE), total lipids (M-LDL-L), total particle concentration (M-LDL-P), phospholipids (M-LDL-PL)<br>For small particles: total cholesterol (S-LDL-C), total lipids (S-LDL-L), total particle concentration (S-LDL-P) |
|  | VLDL subclass particle constituents | Strong | For small particles: total cholesterol (S-VLDL-C),<br>For extra-small particles: total lipids (XS-VLDL-L), phospholipids (XS-VLDL-PL) |
|  | Other | Strong | HbA1c, Apolipoprotein B, total LDL cholesterol, total serum cholesterol |
| Proteins | Chemokine activity | Strong | FAM3B, FAM3D, MIP-5, TECK, |
|  |  | Suggestive | CCL28 |
|  | Chemokine receptors | Strong | IL-3RA, HGF receptor, sGP130, VEGF-R2, VEGF-R3 |
|  |  | None | TCCR |
|  | Receptor function and/or signalling | Strong | F177A, GP116, IGF-1R, IR, JAG1, MBL, PEAR1, PYY, SECTM1, SEMA6A, TLR4 |
|  |  | Suggestive | PLXB2 |
|  |  | None | CD109, CD209, GFRAL, GPIV, LIF-R, Notch-1, PEAR1, sTIE1, sTIE2 |
|  | Cell adhesion | Strong | Cadherin-1, E-selectin, Endoglin, ICAM-4, ISLR2, Laminin, NCAM-L1, OX2G, P-selectin, sICAM-1, sICAM-2, sICAM-5 |
|  |  | None | ADAM23, BCAM, Cadherin-5, Desmoglein-2, ESAM |
|  | Enzyme function | Strong | B3GN2, B4GT1, B4GT2, Cathepsin-S, CLIC5, DPEP2, FA20B, FUT10, GLCE, GNS, IAP, LPH, MA1A2, NDST1, QSOX2, ST4S6, TPST2, XXLT1 |
|  |  | None | ATS13, BGAT, CEL, CHSTB, DYR, MINP1, TLL1 |
|  | Miscellaneous | Strong | C1GLC, CASC4, GOLM1, KIN17, THSD1, TUFT1, |
|  |  | None | Factor VIII, OBP2B |

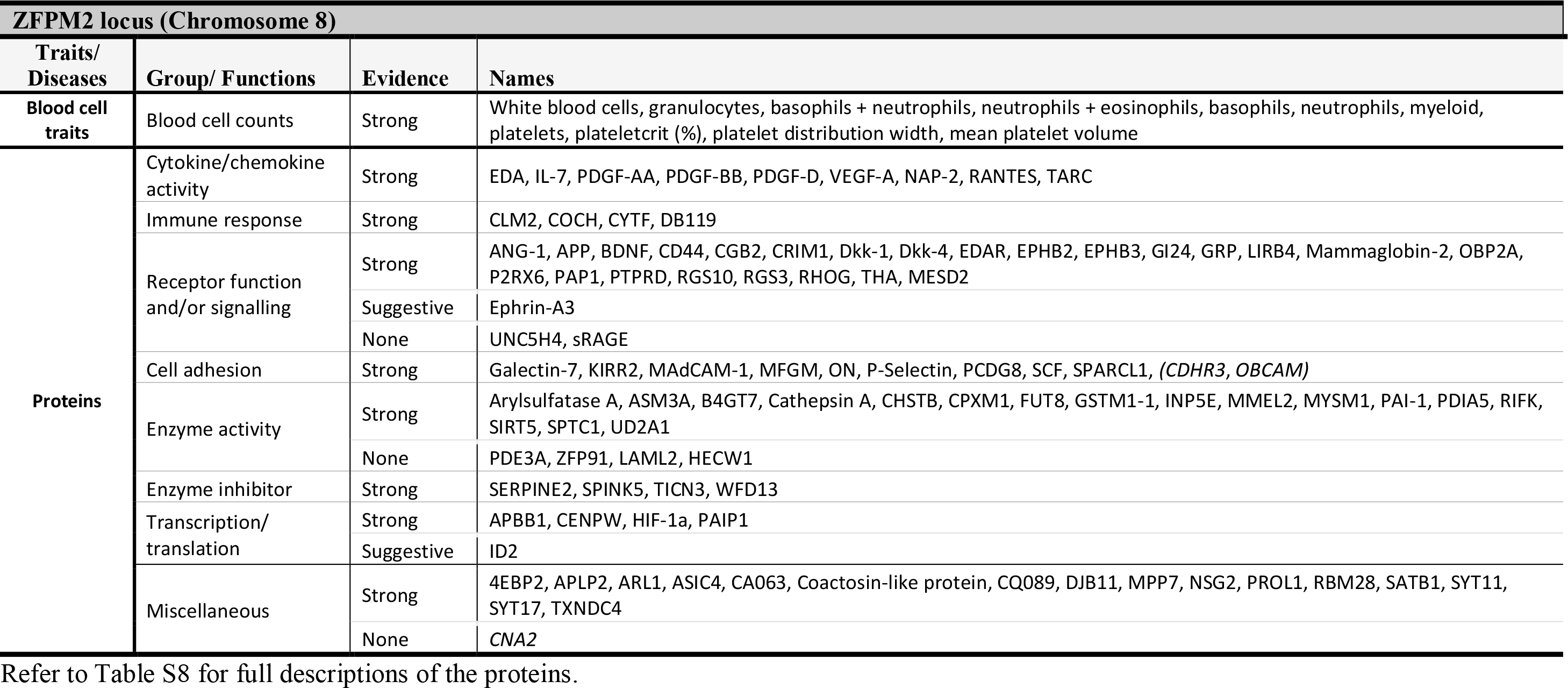

The *ABO* locus colocalised with pQTLs for several membrane proteins (B3GN2, endoglin, GOLM1, OX2G, TPST2) and cell surface receptors (IL-3RA, LIFR, IGF-I R, HGF receptor). *ABO* colocalisation was also observed with pQTLs for adhesion and immune-related molecules involved in leukocyte recruitment, cell adhesion, and transmigration, including sGP130, sICAM-1, sICAM-2, LIRB4, and P-selectin (**Table 3; Table S8**). At the *ZFPM2* locus, colocalisation was seen with pQTLs for proteins generally found in platelet granules (e.g. VEGFA, PDGF-AA, PDGF-BB, PDGF-D, angiopoietin, P-selectin). At the *SERPINE2* locus, we observed that in addition to colocalising with the *cis*-eQTL signal for *SERPINE2* expression, the cytokine network-associated variants colocalised with the *cis*-pQTL variants for SERPINE2 protein levels (**Table S8**). Likewise, the *VEGFA* locus colocalised with a *cis*-pQTL for VEGFA, and the *PDGFRB* locus with a *cis-*pQTL for PDGFRB.

### Relationships of cytokine network variants with complex traits and diseases

Using the NHGRI GWAS Catalog (57,58), we found that, across all eight cytokine network loci, 55 SNPs matched SNPs previously associated with quantitative traits and diseases. (**Table S9**). The lead cytokine network variant at *ZFPM2* (rs6993770) has previously been associated with various platelet traits, including platelet count, distribution width, plateletcrit (total platelet mass) and mean volume (17,59) (**Table S9**).

Next, GWAS summary statistics from a broad range of traits and diseases (**Table S10**), including hematopoietic traits, circulating metabolites, immune- and cardiometabolic-related diseases were compiled for colocalisation analysis with the cytokine network loci. The two cytokine network-associated loci, *ABO* and *ZFPM2*, exhibited strong evidence of colocalisation for several traits and diseases. The *ZFPM2* locus not only colocalised with signals for several platelet trait associations, but also with other haematological trait-associated signals including white blood cell counts, and specifically neutrophil and basophil counts (**Table 3; Table S11**). The *ABO* locus showed colocalisation with various QTLs for haematological traits including red blood cell traits (haemoglobin concentration, red blood cell count, and hematocrit) and white blood cell counts, including granulocyte count and specifically eosinophil count (**Table 3; Table S11**). This is consistent with the *ABO* locus being identified as a pQTL for proteins involved in leukocyte activation as identified previously. Cytokine network variants at the *ABO* locus colocalised with those of intermediate density, low density, and very low-density lipoprotein subclasses as well as glycosylated haemoglobin (HbA1c) (**Table 3; Table S11**), suggesting both inflammatory and metabolic effects. Notably, the same cytokine network variants at the *ABO* locus also strongly colocalised with signals associated with coronary artery disease (CAD), pulmonary embolism, ischemic stroke, and type 2 diabetes (T2D) (**Table 3, Table S11**).

## Discussion

In this study, we first identified a network of 11 correlated cytokines which are known to participate in a broad array of immune responses in circulation. These cytokines include those involved in the classical T_H1_ (IL-12, IFN-γ), T_H2_ (IL-4, IL-6, and IL-10), T_H17_ (IL-6, IL-17, and G-CSF), and T_reg_ (IL-10) responses (51,52) as well as the promotion of angiogenesis, tissue repair and remodelling typically coinciding with inflammatory and post-inflammatory states (VEGF-A, FGF-basic and PDGF-BB) (53). Although previous *in vitro* challenge studies (20,21) indicate antagonistic relationships amongst selected cytokines in the network, our analyses in >9,000 individuals are consistent with previous study utilising similar data (19), showing that these 11 circulating cytokines are positively correlated in the general population. Therefore, at the population level, it is more likely that an equilibrium in circulating levels of disparate cytokines exists, possibly maintained by counter-regulatory mechanisms.

Our multivariate GWAS meta-analysis identified eight loci associated with the cytokine network; confirming six previously-reported associations for circulating cytokine levels (14,16,19) as well as uncovering two additional signals (*PDGFRB* and *ABO)*, empirically demonstrating the statistical power of multivariate approaches. Further, integrative genetic analyses revealed evidence for shared genetic influences between these loci, molecular QTLs, and complex trait and disease associations. This study identified several regions harbouring cytokine-associated signals that colocalise with whole blood and/or immune cell-specific *cis*-eQTLs for a number of genes, including *SERPINE2, ABO*, and *PCSK6*, suggesting these genes are possible candidates underlying the collective expression of cytokines in the cytokine network - or vice versa. Our findings also highlight that the cytokine network associations at the pleiotropic loci, *ABO* and *ZFPM2*, overlap with signals associated with multiple traits, including cardiometabolic diseases, immune-related proteins, and platelet traits.

*SERPINE2* encodes protease nexin-1, an inhibitor of serine proteases such as thrombin and plasmin, and is therefore implicated in coagulation, fibrinolysis and tissue remodelling (60). It shares similar functions with its better-known homolog *SERPINE1*, or plasminogen activator inhibitor-1 (PAI-1), the elevation of which is associated with thrombosis and cardiovascular risk (60). However, there is also evidence that *SERPINE2* has pleiotropic roles in immune and inflammatory regulation, that could be either dependent or independent of its function as a serine protease. It is expressed in many tissue types, and its expression can be induced by pro-inflammatory cytokines such as IL-1α (61,62). Conversely, *SERPINE2* can itself influence inflammatory status: *SERPINE2* is a candidate susceptibility gene for chronic obstructive pulmonary disease, and *SERPINE2*-knock-out mice exhibited extensive accumulation of lymphocytes in the lungs, through a mechanism linked to thrombin and NFκB activation (62). We observed in our data that the cytokine network associations overlapped with the *SERPINE2* pQTL signal. Moreover, using immune cell-specific *cis*-eQTL data, we further demonstrated colocalisation between the cytokine network and *SERPINE2 cis*-eQTL signals specifically in CD4^+^ T-cells and B-cells. This suggests that the association between *SERPINE2* and the cytokine network at this locus is at least partially-driven by lymphocytic expression - consistent with *SERPINE2* itself influencing chemotaxis and recruitment of lymphocytes (62). Our analyses demonstrate that the importance of *SERPINE2* in regulating immune and inflammatory processes is potentially greater than previously anticipated, and warrants further targeted research.

Like *SERPINE2*, the *ABO* locus has widespread pleiotropic effects. The most well-known function of *ABO* is its determination of blood group. The human *ABO* gene has three major alleles (A, B, and O) that determine ABO blood type. The A and B alleles encode for distinct “A” versus “B” glycosyltransferases that add specific sugar residues to a precursor molecule (H antigen) to form A versus B antigens, respectively (63). The O allele results in a protein without glycosyltransferase activity (63). The lead cytokine-associated variant rs550057 and its proxies in moderate LD (r^2^ = 0.6; rs507666, rs687289) have been previously shown to determine the *ABO* allele (64), but they have also been associated with circulating levels of inflammatory proteins such sICAM-1, P-selectin, and ALP (17,65,66). Our study showed that cytokine network associations at the *ABO* locus share colocalised signals with a host of other proteins and traits, including lipoproteins (IDL, LDL, VLDL), proteins of immune function, immune cell subsets, and cardiometabolic diseases (**Table 3**), highlighting the potential for shared molecular etiology amongst these traits. Our analyses highlight the potential genetic basis for numerous previous observations linking ABO blood group to an array of similar traits and phenotypes (18,67–71).

It could therefore be speculated that the *ABO* gene influences the risk of cardiometabolic disease due to its involvement in multiple inflammatory, haemostatic and metabolic processes; however, our current understanding of the mechanisms behind this remains unclear. For instance, non-O blood groups have been associated with increased risk of both cardiovascular disease, venous thromboembolism, stroke, and T2D (68,72). However, the O blood group has itself been linked to elevated IL-10 and worse outcomes given existing coronary disease (risk of cardiovascular death, recurrent myocardial infarction and all-cause mortality) (64). Other studies have suggested a role for von Willebrand factor (VWF), a coagulative factor which also expresses ABO antigens - in particular, the O phenotype is associated with lower VWF, which may explain reduced thrombotic and cardiovascular risk (64,73). It has been suggested that the link between ABO blood group type and venous thromboembolism (VTE) is potentially driven by VWF and Factor VIII - non-O blood group individuals presented a higher risk of venous thromboembolism and had elevated levels of both VWF and Factor VIII (74,75). Also relevant is the link between *ABO* and adhesion molecules such as E-selectin and sICAM-1 which are overexpressed in inflammatory states (18,66,70,71). sICAM-1 is a known positive correlate with cardiovascular disease; however, it is the A blood group, not O, that is associated with reduced sICAM-1 levels, again complicating the picture (70). Inferring the exact causal relationships amongst all these entities will require intricate follow-up experimental investigation, involving simultaneous examination of all key players. It is particularly unclear whether the link with cardiometabolic diseases may be due to its direct modification of H antigen, or on the glycosyltransferase activity of the encoded enzyme on other proteins, or some combination of both. In our study, formal causal inference (e.g. with Mendelian Randomisation) was not possible because the corresponding multivariate beta-coefficients and standard errors are not currently calculable and the locus itself has extensive pleiotropy.

The *ZFPM2* locus has been associated with platelet traits (59), and our findings highlight its importance as a determinant of platelet and angiogenic cytokine activity. *ZFPM2* encodes a zinc finger cofactor that regulates the activity of GATA4, a transcription factor reported to play a critical function not only in heart development (76) but also modulation of angiogenesis. In particular, GATA4 directly binds to the promoter of angiogenic factor *VEGFA* and regulates its expression (77), and it has been shown that disruption of ZFPM2-GATA4 interaction alters the expression of *VEGFA* and other angiogenesis-related genes (78). VEGFA and PDGFR-BB, which are part of the cytokine network, have been found to be released via alpha granules of activated platelets, and serum VEGFA levels correlate closely with blood platelet counts (79–81). In our study, we show that the cytokine-associated signal at the *ZFPM2* locus colocalised with GWAS signals for platelet traits and platelet proteins. The lead cytokine network SNP rs6993770 has been reported to be a *trans*-eQTL in whole blood for gene products typically found in platelets and their receptors (e.g. CXCL5, GP9, MYL9, VWF) (46). Collectively, these findings suggest that this locus regulates the number and/or cytokine activity of circulating platelets, and that this potentially occurs via interaction with *GATA4* and regulation of *VEGFA*.

In conclusion, our study illustrates the utility of multivariate analysis of correlated immune traits and highlights potentially fruitful avenues of biological investigation for multivariate genetic signals. Our results highlight that certain gene loci drive the expression of a cytokine network with immune, inflammatory and tissue repair functions; and, simultaneously, these loci are implicated in the regulation of other haemostatic and metabolic functions, with relevance to human health and disease. This stresses the fact that the processes of inflammation, haemostasis and repair often run concurrent with each other after injury, and that biological systems often feature ample redundancy and feedback loops within individual effectors.

## Acknowledgements

Artika Nath was supported by an Australian Postgraduate Award. This research was supported in part by the Victorian Government’s OIS Program. Michael Inouye was supported by an NHMRC and Australian Heart Foundation Career Development Fellowship (no. 1061435). Gad Abraham was supported by an NHMRC Early Career Fellowship (no. 1090462). Qin Qin Huang is supported by the Melbourne International Research Scholarship. The Young Finns Study has been financially supported by the Academy of Finland: grants 286284, 134309 (Eye), 126925, 121584, 124282, 129378 (Salve), 117787 (Gendi), and 41071 (Skidi); the Social Insurance Institution of Finland; Competitive State Research Financing of the Expert Responsibility area of Kuopio, Tampere and Turku University Hospitals (grant X51001); Juho Vainio Foundation; Paavo Nurmi Foundation; Finnish Foundation for Cardiovascular Research; Finnish Cultural Foundation; The Sigrid Juselius Foundation; Tampere Tuberculosis Foundation; Emil Aaltonen Foundation; Yrjö Jahnsson Foundation; Signe and Ane Gyllenberg Foundation; Diabetes Research Foundation of Finnish Diabetes Association; and EU Horizon 2020 (grant 755320 for TAXINOMISIS); and European Research Council (grant 742927 for MULTIEPIGEN project); Tampere University Hospital Supporting Foundation. Peter Würtz is supported by the Novo Nordisk Foundation (15998) and Academy of Finland (312476 and 312477).

