## Supplemental Figures for "Multivariate genome-wide association analysis of a cytokine network reveals variants with widespread immune, haematological and cardiometabolic pleiotropy"

### Supplementary Text

### Figures

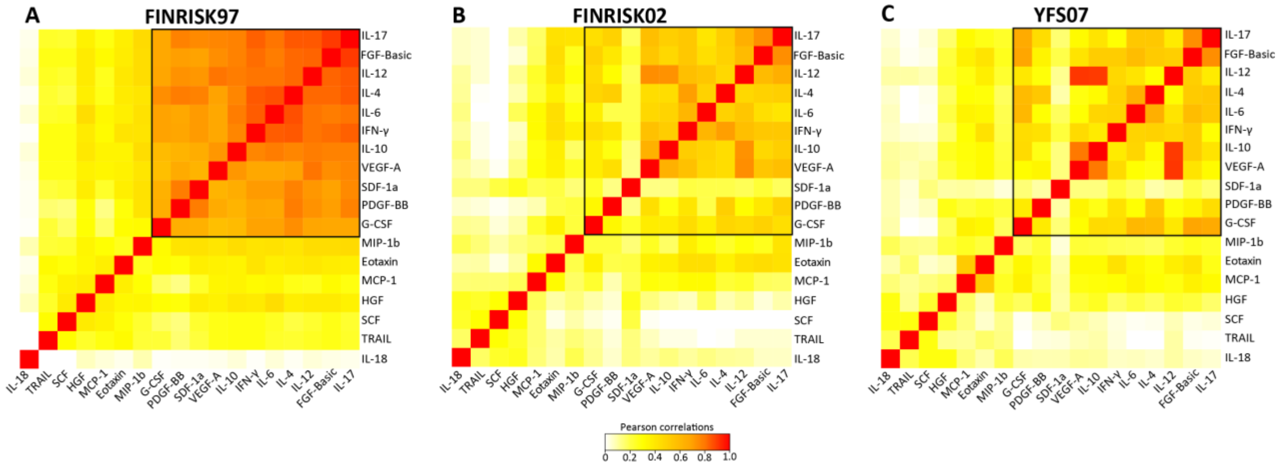


**Figure S1: Comparison of cytokine-cytokine correlation in FINRISK07, FINRISK02, and YFS07.**

The heatmaps show the correlations between the normalised cytokines residuals in the discovery dataset, **(A)** FINRISK97, and the replication datasets, **(B)** FINRISK02 and **(C)** YFS07. Each square represents the Pearson's correlation coefficient between the cytokines. The black box shows the correlation patterns among the 11 correlated cytokines (discovered using the FINRISK97) across the three datasets. The correlation matrix in FINRISK07 was subjected to hierarchical clustering using distance as 1 minus the absolute value of the correlations. The ordering of rows and columns in FINRISK02 and YFS07 was defined by the ordering in FINRISK07. The strength of the correlations is indicated by the colour on the scale, where red is a high positive correlation.

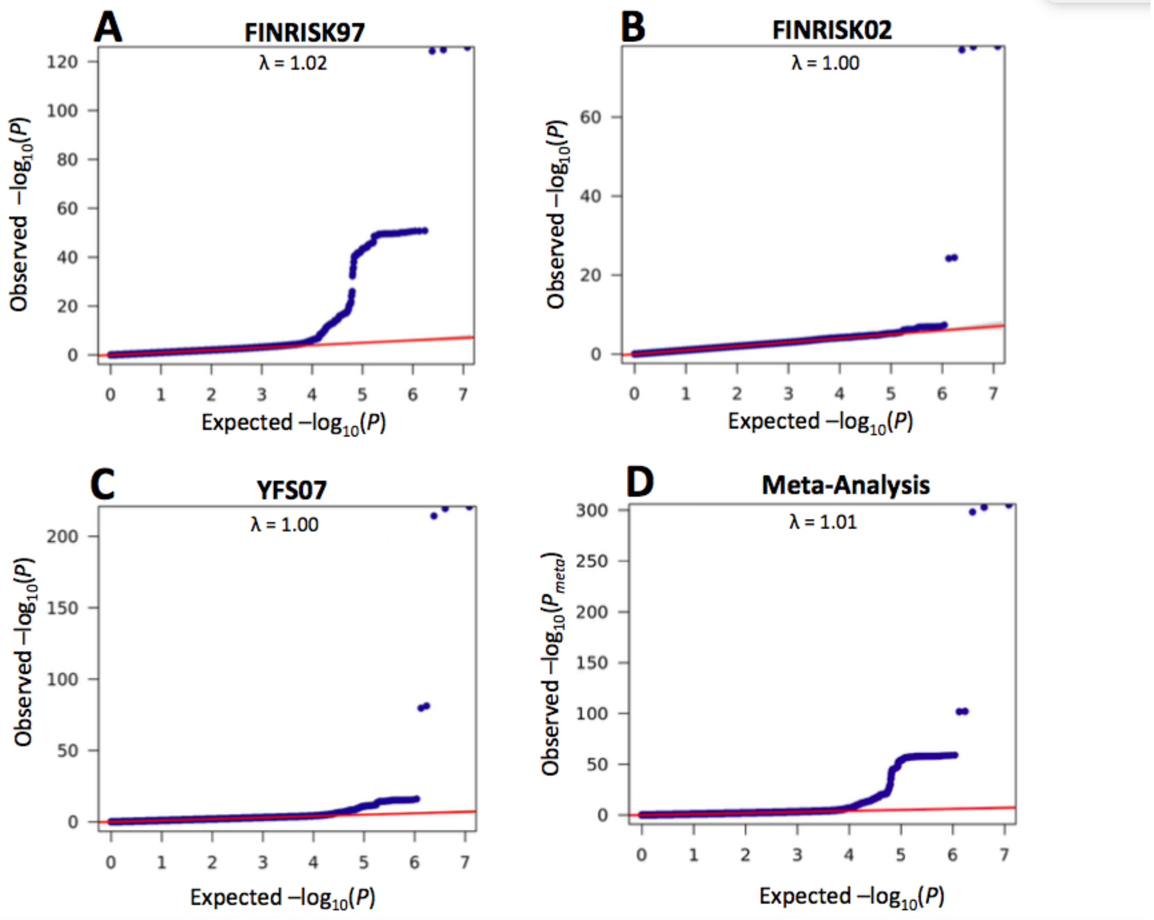


**Figure S2: Quantile-quantile (Q-Q) plots resulting from the multivariate GWAS in the three cohorts separately and meta-analysis.**

Q-Q plots of observed (*y*-axis) *vs*. expected *P* values (*x*-axis) for each SNP from the multivariate genome-wide association in **(A)** FINRISK97, **(B)** FINRISK02, **(C)** YFS07, and **(D)** Meta-analysis of the three cohorts. The diagonal red line (*y*=*x*) indicates null hypothesis of no association. The inflation factor (λ) was between 1.0 – 1.02 suggesting that inflation from population substructure or other confounders was appropriately adjusted for.

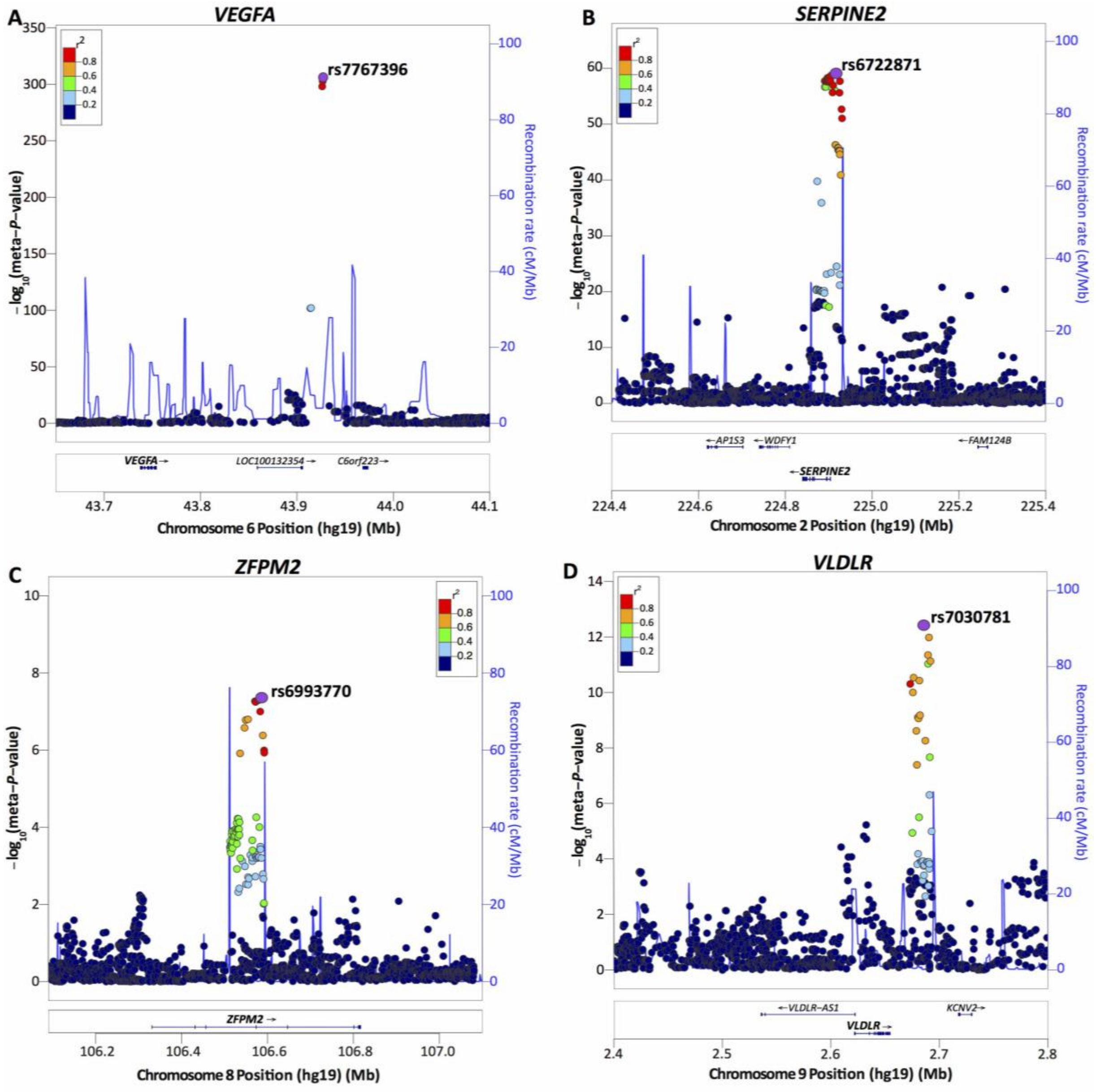


**Figure S3: Regional association plots for each of the 8 loci associated with the cytokine network from the meta-analysed multivariate GWA analysis.**

**(A)** ***VEGFA*** **locus**, rs7767396 is an intergenic SNP located 172.83kb downstream of vascular endothelial growth factor A (*VEGFA*) gene on chromosome 6p21.1. **(B)** ***SERPINE2*** **locus**, rs6722871 lies 10.9kb upstream of *SERPINE2* on chromosome 2q36.1. **(C)** ***ZFPM2*** **locus**, rs6993770 lies within intron 4 of the zinc finger protein multitype 2 (*ZFPM2*) gene on chromosome 8q23.1. **(D)** ***VLDLR*** **locus**, rs7030781 is situated ~31.8kb away from the very low-density lipoprotein receptor (*VLDLR*) gene on chromosome 9p24.2. For each plot, the circles represent the -$\log_{10}$ meta-analysed *P*-values (*y*-axis) of SNPs plotted against their chromosomal position (*x*-axis). The lead SNP in each plot is denoted by a purple circle, and its pairwise LD ($r^2$) strength with other SNPs in the region, estimated from the “1000 genomes Mar 2012 EUR” population, is indicated by color. The blue lines indicate the recombination rates. The plots were generated using the LocusZoom online tool (http://locuszoom.sph.umich.edu/locuszoom/).

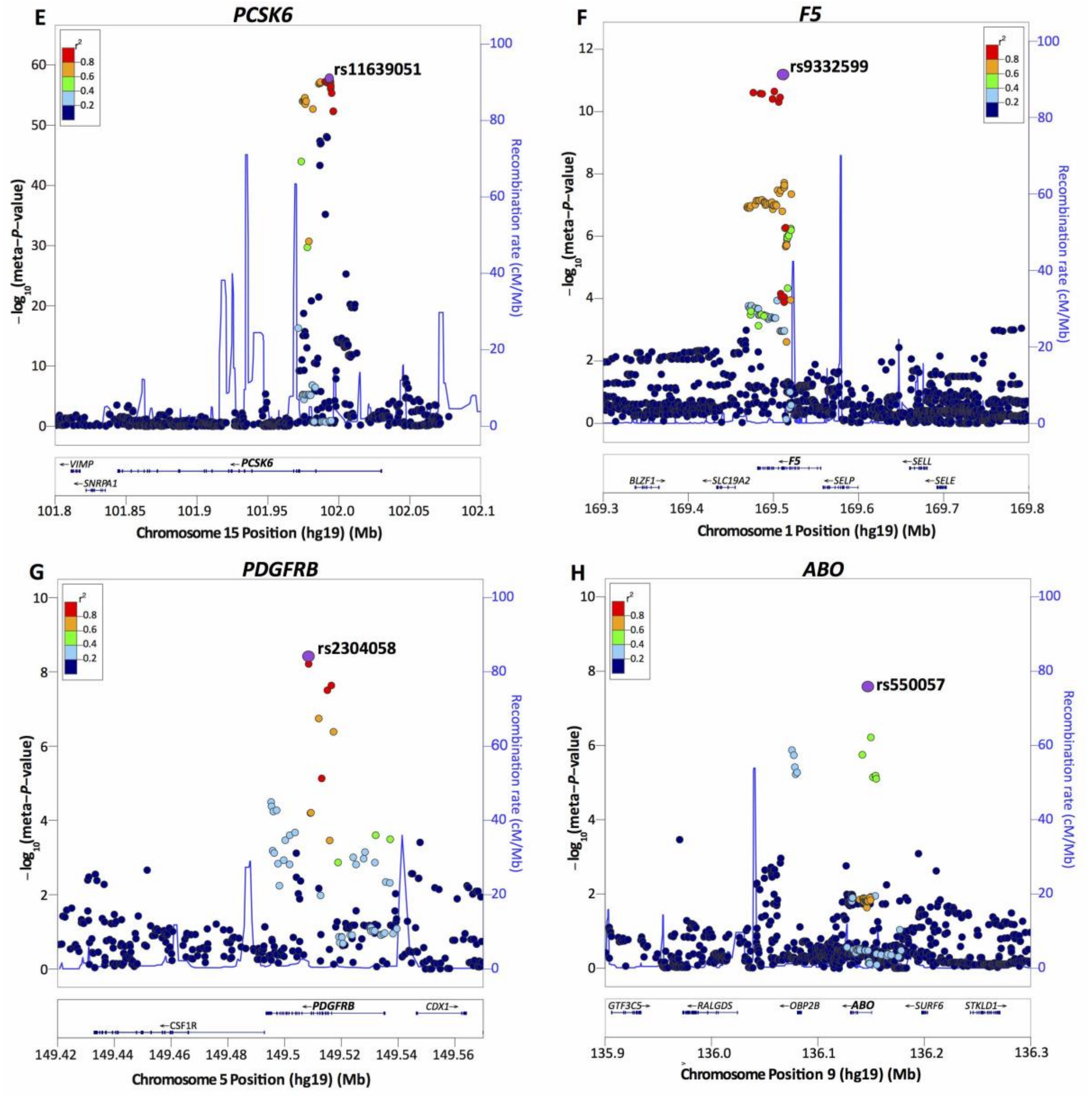


**Figure S3: Regional association plots for each of the 8 loci associated with the cytokine network from the meta-analysed multivariate GWA analysis**

**(E) *PCSK6* locus**, rs11639051 is located in the second intron of *PCSK6* (proprotein convertase subtilisin/kexin type 6) on chromosome 15q26.3. **(F) *F5* locus**, rs9332599 is located within intron twelve of factor V (*F5*) gene on chromosome 1q24.2. **(G) *PDGFRB* locus**, rs2304058 lies within the tenth intron of the platelet-derived growth factor receptor-beta (*PDGFRB*) gene on chromosome 5q32. **(H) *ABO* locus**, rs550057 is located within the first intron of *ABO* gene on chromosome 9q34.2.

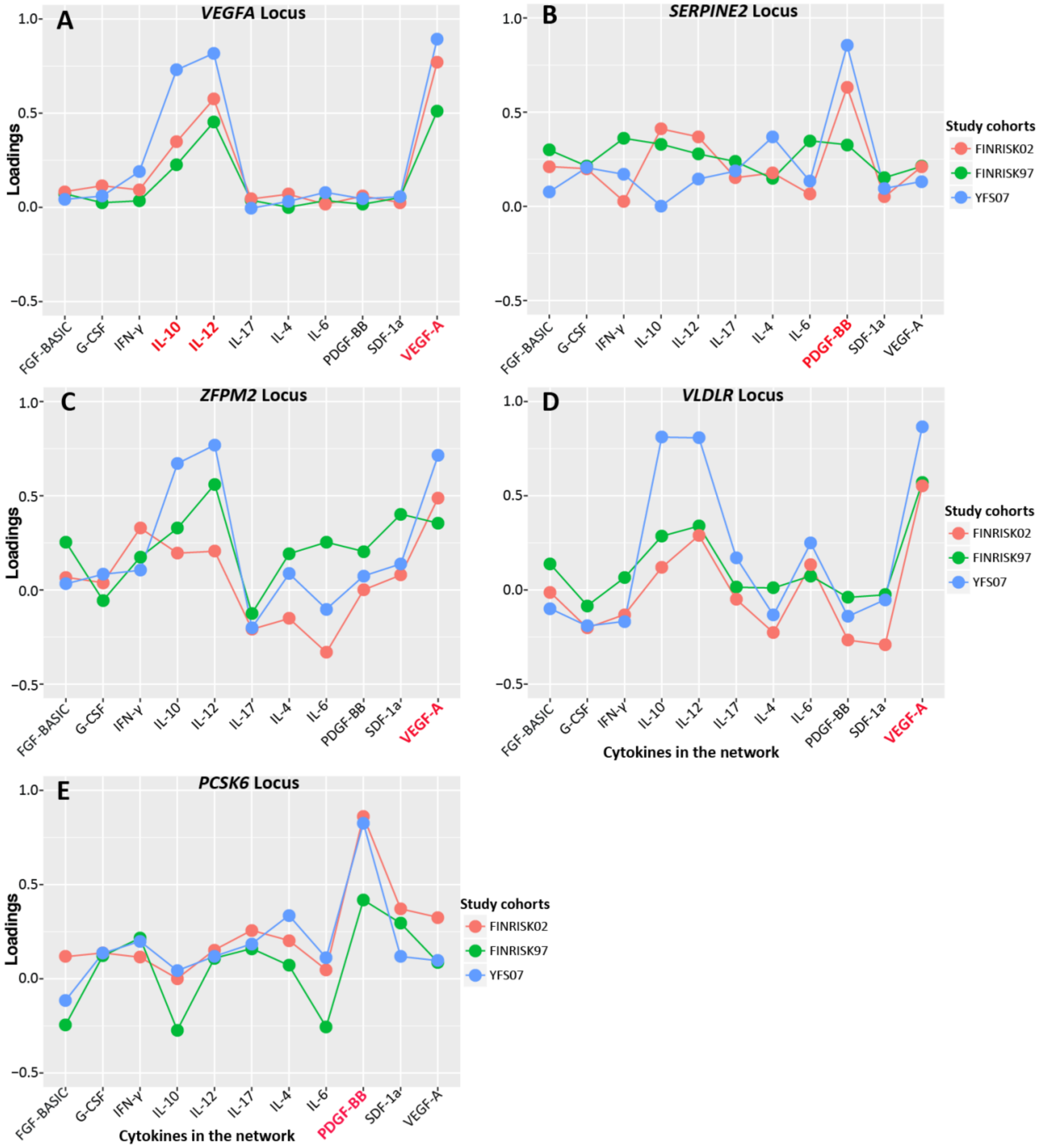


**Figure S4: Loadings, the contribution of each cytokine in the cytokine network to the multivariate association results with the lead SNPs at the *VEGFA*, *SERPINE2*, *ZFPM2*, *VLDLR* and *PCSK* locus in each study cohort.**

The trait loadings are output results from MV-PLINK – the sign of the loadings for each cytokine in each cohort indicates whether the genetic variant influences different cytokines in the same or opposite effect direction.
